# Oculomotor dance learning task: Implications for audio-visual cued spatial learning

**DOI:** 10.64898/2026.01.23.701340

**Authors:** Michael Petrovski, Susu Beheiry, Udichi U. Das, Simran Rooprai, Ashkan Karimi, Jenny R. Simon, Rachel J. Bar, Joseph F. X. DeSouza

## Abstract

Motor sequence learning involves the coordination of both oculomotor and manual motor systems through the practiced repetition of a fixed sequence of actions. This study aims to address whether a sequence-based learning paradigm centered on the visual-motor system can feasibly be measured while listening to music (Bar and DeSouza 2016). It aims to develop a new visual-motor-based learning paradigm with music, potentially promoting neuroplasticity and creating new interventional tools, building upon prior research that shows behavioural and putative neural changes following dance-based neurorehabilitation in people with Parkinson’s disease (Bearss et al. 2024). Eye movements of 10 participants (8 female, 2 male) were tracked using the Eyelink 1000 Plus system during a 68-second eye-dance sequence. The experiment consisted of a learning phase, where participants observed the sequence five times with 30-second breaks, and a performance phase, where they performed the sequence five times from memory on a grey screen without visual cues. Music was incorporated into both phases to aid memorization of the 4 spatial locations. After each performance, the participant was shown a visual reinforcer and asked for their thoughts on how well they executed the dance. A visual reinforcer flashes one of three different colours: red, yellow, or green. Each colour corresponds to how many steps in the dance a participant performed correctly, with key points being: under one third, between one to two thirds, and over two thirds of total steps correct. Participants were scored based on timing of the steps as well for exact (1.00), good (0.66), slightly off (0.33) or missed (0) steps. Data was analyzed using R4.3.1, MATLAB, and Experiment Builder: Data Viewer software. Results showed a significant improvement in performance accuracy between the first session (g1; M = 40%, SD = 7.2%) and the last session (g5; M = 69.7%, SD = 22.8%). A repeated-measures ANOVA revealed a significant main effect of session on performance accuracy, F(4, 36) = 6.99, p < 0.001, η²_G_ = 0.26, indicating that accuracy significantly improved over sessions. Post-hoc Bonferroni comparisons showed that accuracy in later sessions was significantly higher than earlier sessions, suggesting a defined learning curve and consolidation of performance pattern across repeated practice. Similarly, there was significant improvement in timing accuracy between the first session g1; M = 0.29, SD = 0.06) and the fifth session (g5; M = 0.46, SD = 0.12). A repeated-measures ANOVA revealed a significant main effect of session on timing precision, F(4, 36) = 11.67, p < 0.001, η²_G_ = 0.25, indicating significant improvements in temporal control and coordination over sessions. Post-hoc Bonferroni comparisons showed that timing precision significantly improved between early and late sessions (e.g, g1-g4, p <0.01; g1-g5, p < 0.001), suggesting a defined learning curve and increase in precision across repeated practice. These findings suggest that visual-motor-based interventions have the potential to enhance motor and non-motor symptoms like depression and anxiety for neurodegenerative diseases such as Parkinson’s Disease (PD). The results provide a foundation for developing targeted therapies that integrate learning paradigms to improve functional outcomes, warranting further exploration of their long-term efficacy.

## Introduction

Motor sequence learning involves the coordination of both oculomotor and manual motor systems through the practiced repetition of a fixed sequence of actions, resulting in automatized execution of movement (Rubino et al., 2025). While extensive research regarding motor sequence learning has been explored involving neural bases of behaviour (Jäger et al., 2022, Tremblay et al., 2021), the oculomotor context remains an emerging field of motor sequence learning. Typical design of motor sequence learning tasks generally assess reaction times using limb movement in response to stimuli (Jeyarajan et al., 2024; Maceira-Elvira et al., 2022; Rubino et al., 2025; Sailer et al., 2005; Vieluf et al., 2015). Although within a smaller subset of motor sequence learning literature, various studies found that eye movements predict motor sequence learning similar to that of physical movement (Coiner et al., 2019; Medimorec et al., 2021; Rubino et al., 2024), and have activational patterns closely resembling physical motor response (Rubino et al., 2024; Rubino et al., 2025; Sailer et al., 2005). Other studies have found auditory stimuli to enhance motor sequence learning (Leow et al., 2025), aiding response to visual stimuli. The use of auditory cues synchronously within sequence learning has been associated with increased development of explicit knowledge of learned sequences (Han et al., 2024). In addition, cues of this nature are found to aid in movement sequences that occur over longer periods of time (Bar & DeSouza, 2016; Leow et al., 2025). Understanding the components behind sequence learning that reinforce behaviours and show greater results in combination rather than individually creates a significant demand for future task design to seamlessly incorporate both facets.

Defining the underlying mechanisms of motor sequence learning, specifically the neurological communications between areas discussed prior, is increasingly critical. Neuroplasticity is the brain’s ability to reorganize neural pathways in response to degeneration or loss of function within such pathways. To prompt neuroplastic change is to engage brain function of specified regions of interest with a task and to understand the compensations or new connections made. Various methods of quantifying neuroplastic changes are documented (Barnstaple et al., 2021; Ferreira et al., 2025; Herzberg et al., 2024; Hänggi et al., 2010; Hehl et al., 2025; Tremblay et al., 2021), however the developments brought about by physical intervention or tasks related to movement will be pertinent to the following body of work. Systematic reviews have highlighted the importance of physical exercise or engagement in improving executive function and wellbeing in old age (Cheng et al., 2024; Du et al., 2025; Quan et al., 2025). Even in tasks involving minimal physical effort, greater activation of regions involved such as frontal eye fields (FEF) (Gonzalez et al., 2016), superior colliculus (Soetedjo et al., 2009), cerebellar oculomotor vermis (Kurkin et al., 2014), and SMA-related motor networks have been shown (Bonzano et al., 2015; Coiner et al., 2019; Rubino et al., 2024; Rubino et al., 2025). In addition, it has been noted that eye tracking proves to be a promising tool for assessment and clinical applications as they can potentially be applied towards measurement of neuroplastic changes in neurodegenerative diseases (Culicetto et al., 2025; Diotaiuti et al., 2025). Providing analysis of specific areas involved in neuroplastic change brought about through motor engagement, whether ocular or physical, remains pivotal.

As a form of physical movement combined with audio influence, and common amongst all cultures and ages, dance has great quantifiable effect on motor sequence learning and neuroplasticity (Gao et al., 2025; Jeyarajan et al., 2024; Johansson et al., 2022; Tsai et al., 2024). Dance actively engages an individual physically, cognitively, and sensorially, and it has been widely recognized as a practical intervention to promote neuroplasticity and execute motor sequence learning (Barnstaple et al., 2021; Cheng et al., 2024; Dhami et al., 2015; Du et al., 2025; Simon et al., 2024; Volpe et al., 2025). While a vast amount of literature involving measures of neuroplasticity within the last decade suggests physical exercise or dance can lead to improvements or mitigation of impairments, the need for both simplified and accessible tasks is paramount. The physical demands of exercise and other coordinated movement such as dance continue to pose the challenge of being inaccessible to various groups. It has been widely urged within the surrounding literature to create simplified versions of both dance intervention (Dhami et al., 2015; Du et al., 2025), and physical exercise intervention (Johansson et al., 2022). For old-aged, neurodegenerative disordered, or individuals with disabilities, severity of cognitive or physical decline is a key issue (Gorges et al., 2016; Hill-Briggs et al., 2007), and it is acknowledged that interventions should come to address individuals beyond the mild-moderate scale.

The aim with the present eye tracking task is to develop a feasible, non-exertive task that minimizes the physical response needed from a participant whilst still activating similar regions of the brain to that of coordinated physical movement and auditory cues such as dance. Within the current study, the usage of eye movement as an alternative to physical movement in motor sequence learning was investigated. In combination with auditory cues of a musical piece, the additional goal of diversifying current literature on multisensory sequence learning was sought. We hypothesized that a within-subjects design with 5 sessions of eye dance practice over 5 weeks of time would result in improvements within sequence-specific learning, namely performance accuracy and timing precision.

## Materials and Methods

Ten healthy adults were recruited from the student population at York University, all of whom provided written informed consent. Exclusion criteria for the study were visual or auditory impairment. Total n = 10 (female = 8, male = 2), Ages: 20-25, M = 22.

Stimuli were displayed on a built-in LED iMac monitor (1680 x 1050 pixels). A headrest was set to ensure stability of participant gaze to the stimuli and clear tracking of the eye. Viewing distance of the display to headrest was measured to 55 cm. The headrest was adjusted to ensure gaze was matched to the direct center of screen. Through training trials, a gray background was presented with a 848 x 480 pixel video centered within the display of a woman directing eye movements of the overall dance pattern. Runtime of the video and accompanying audio piece is 68 seconds. Through performance trials, a gray background was presented with 5 red outlined squares each measuring 200 x 200 pixels. The squares were located on the display and labelled as the following positions: Center (840, 525), Up (840, 100), Down (840, 950), Left (100, 525), and Right (1580, 525). All eye-movements were directionally cued along the X and Y axes recorded within an eye-tracking paradigm on an Eyelink 1000 plus (EYELINK I CL v5.04 Sep 25 2014 CAMERA: Eyelink GL Version 1.2 Sensor=AJ7).

Participants were taught a complex eye movement sequence to a 68 second musical piece by watching a choreographed eye-dance pattern. The video and music of the dance were played five times with thirty second break intervals featuring a gray interface with no audio in between. The set of prior events is referred to as the learning phase of the test. Following the learning phase, participants are tasked to perform the same sequence of eye movements using only the musical piece and target markers on a gray interface. Executions of the dance sequence were recorded five times with a thirty second break with a gray interface similar to the learning phase and are referred to as the performance phase. At the end of each 68 second performance recording, 5 out of the 30 seconds within the break consisted of a colour flash of the interface corresponding to the accuracy of steps correctly made. Three colours were shown: red, yellow, and green; corresponding to total score being <33% correct, 33%< x <66% correct, and >66% respectively. The visual reinforcer was shown to cue accuracy of the pattern recorded of each participant. Participants were also asked to rate their performance after each trial on a 1-100% scale. After learning and performance phases the session was completed, participants were instructed to complete their next session within 7-10 days.

Data plotting and analysis were conducted using softwares R 4.3.1 and Matlab (R2024b, MathWorks). Results of accuracy statistics were found using a repeated-measures ANOVA. Eye gaze data was sampled every 2ms, all blinks and positions off-screen were omitted from data processing. Target thresholds were located within 200 pixel squares around exact targets (Fig. 2). Movements were only marked if 200 samples remained within target threshold, then compared with the dance sequence to be matched. One-way repeated-measures ANOVAs were selected as the statistical criteria of both performance and timing accuracy to assess learning over time. Both analyses were performed with 95% confidence intervals and significance threshold at *p* ≤ 0.05.

**Fig 1.**
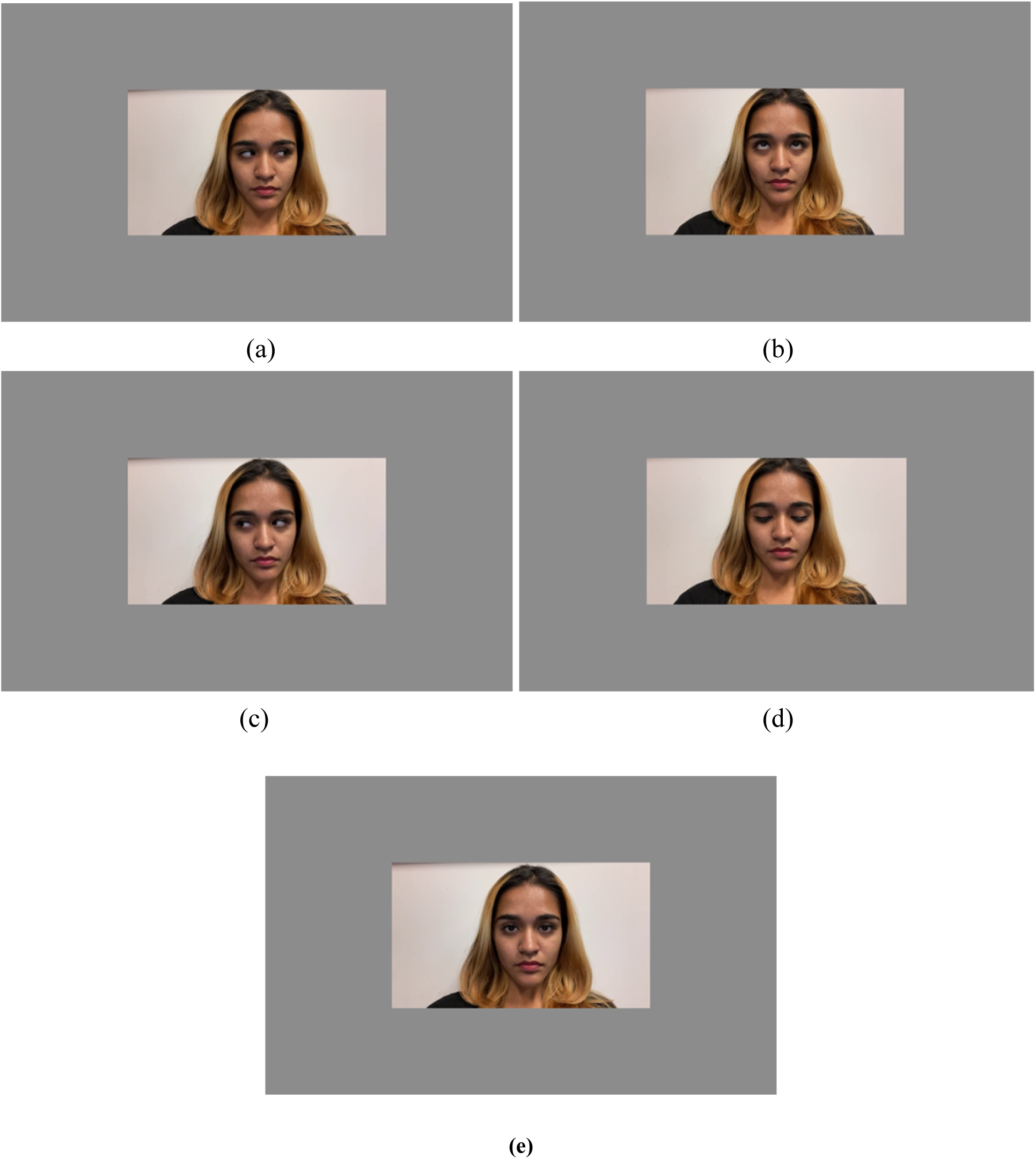
Training Interface. Interface was shown to each participant for 68 seconds with accompanying music and choreography that was instructed to be followed with eye movements. Movements were directionally cued as (a) right, (b) up, (c) left, (d) down, and (e) center.

**Fig 2.**
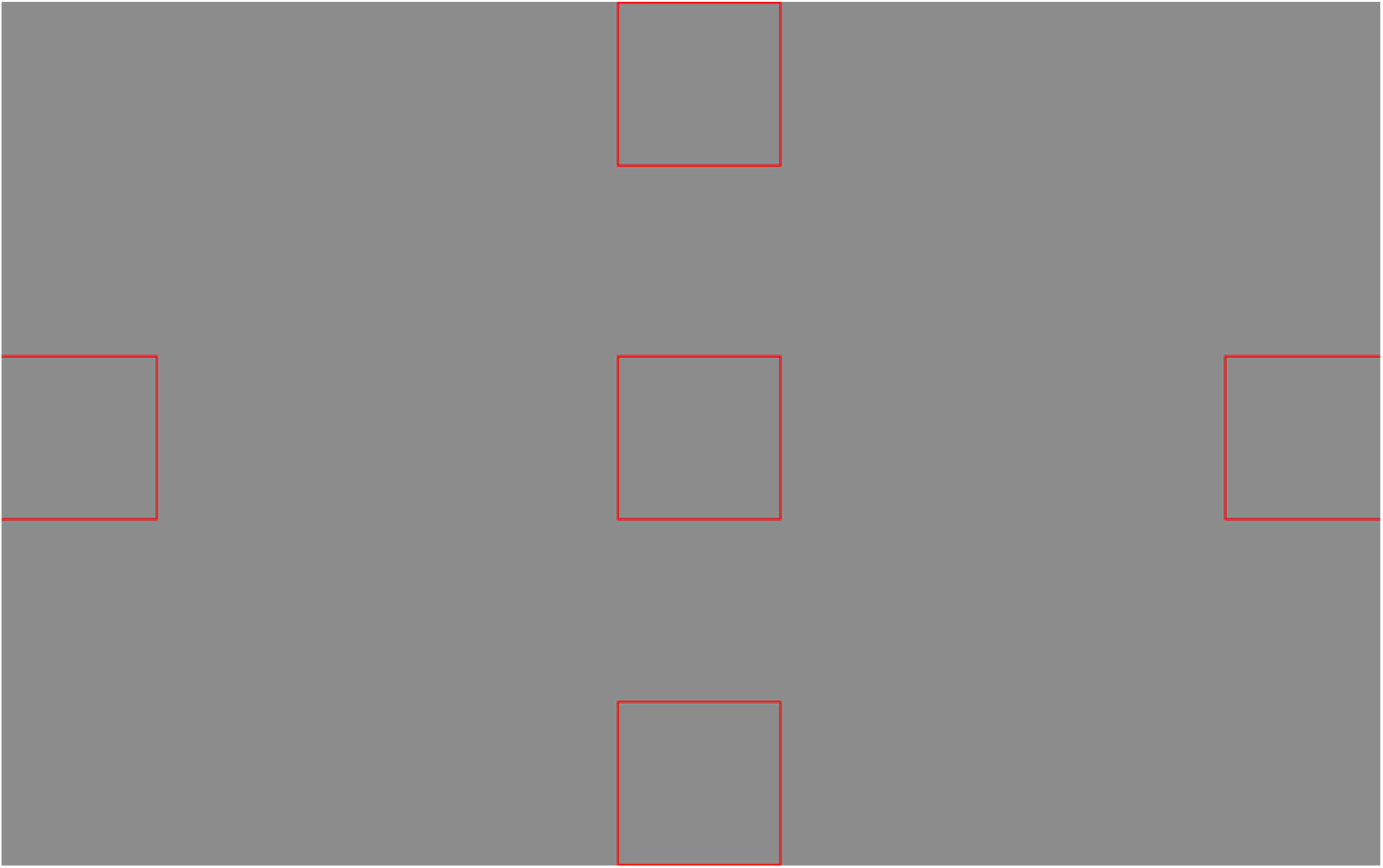
Live Scoring Interface. Interface was shown to each participant for 68 seconds with accompanying music as played in training trials. The red boxes drawn coordinate with the eye movements denoted in Fig 1.

**Fig 3.**
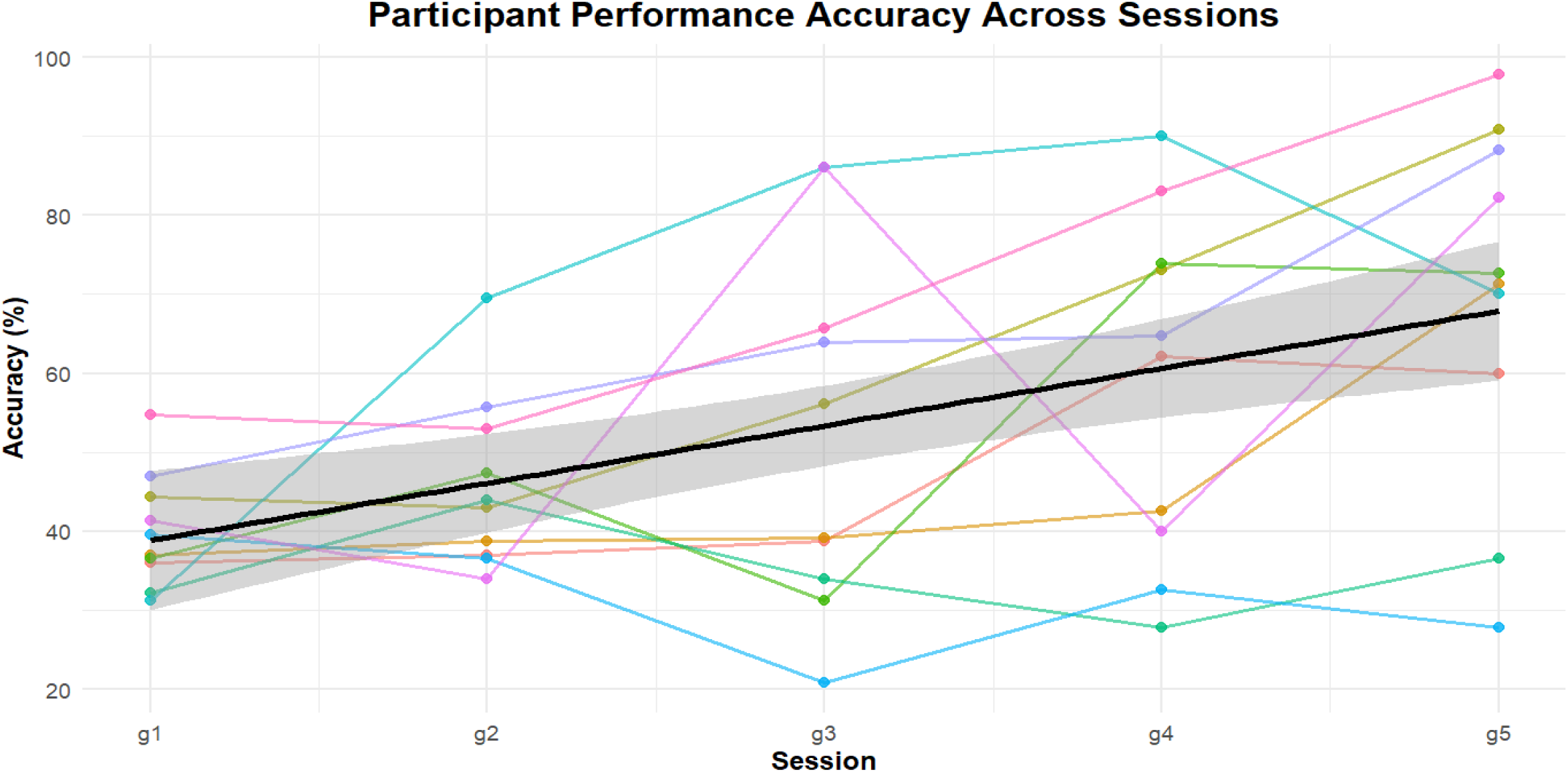
Participant Performance Accuracy Across Sessions. Accuracy was recorded based on the proportion of movements correctly executed to the dance sequence (X/46). A line of best fit and shaded 95% confidence interval is shown as well.

**Fig 4.**
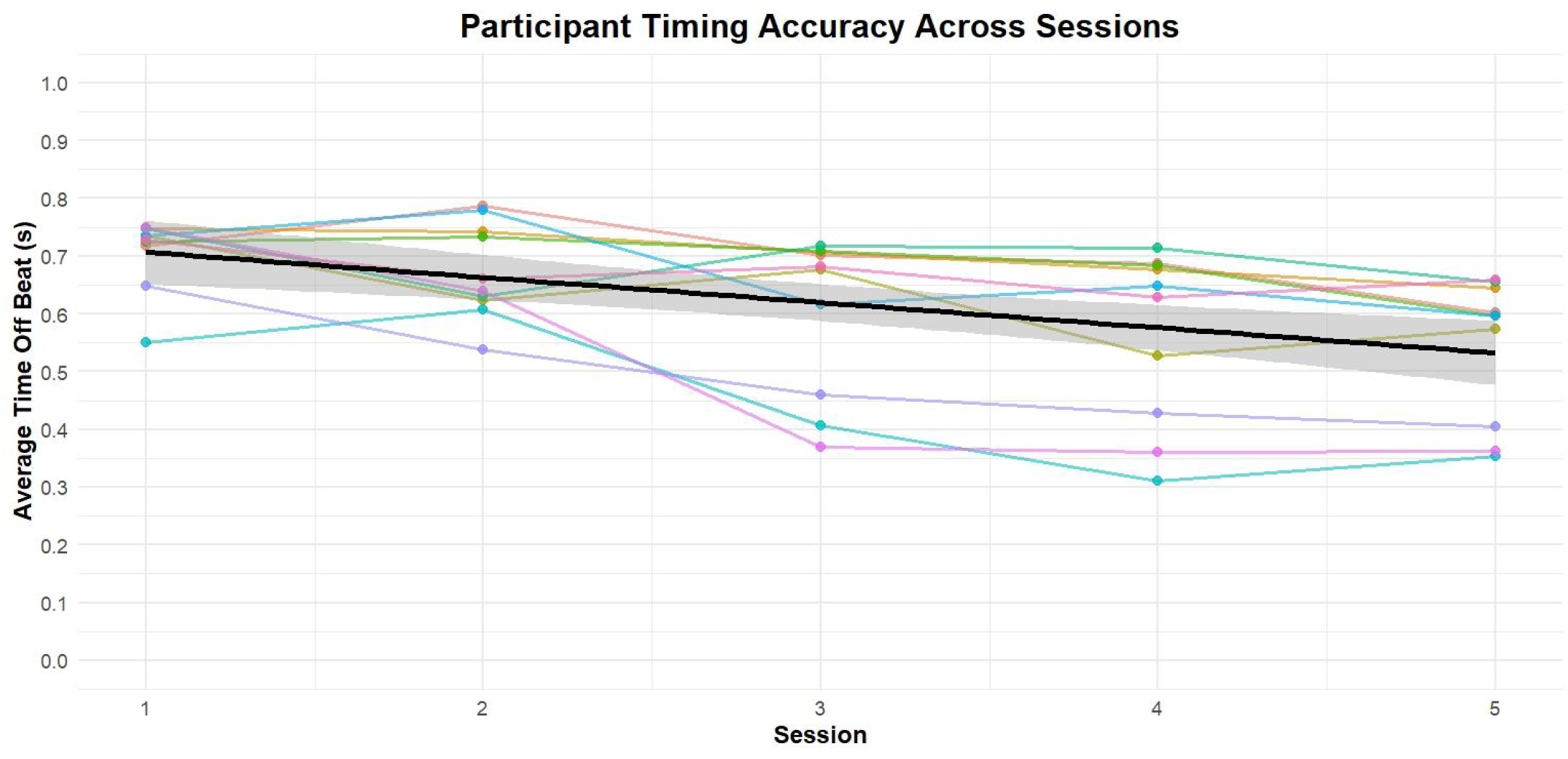
Participant Timing Precision Across Sessions. Participant timing was measured based on proximity to the movements synced to video runtime.. A line of best fit and shaded 95% confidence interval is shown as well.

**Fig 5.**
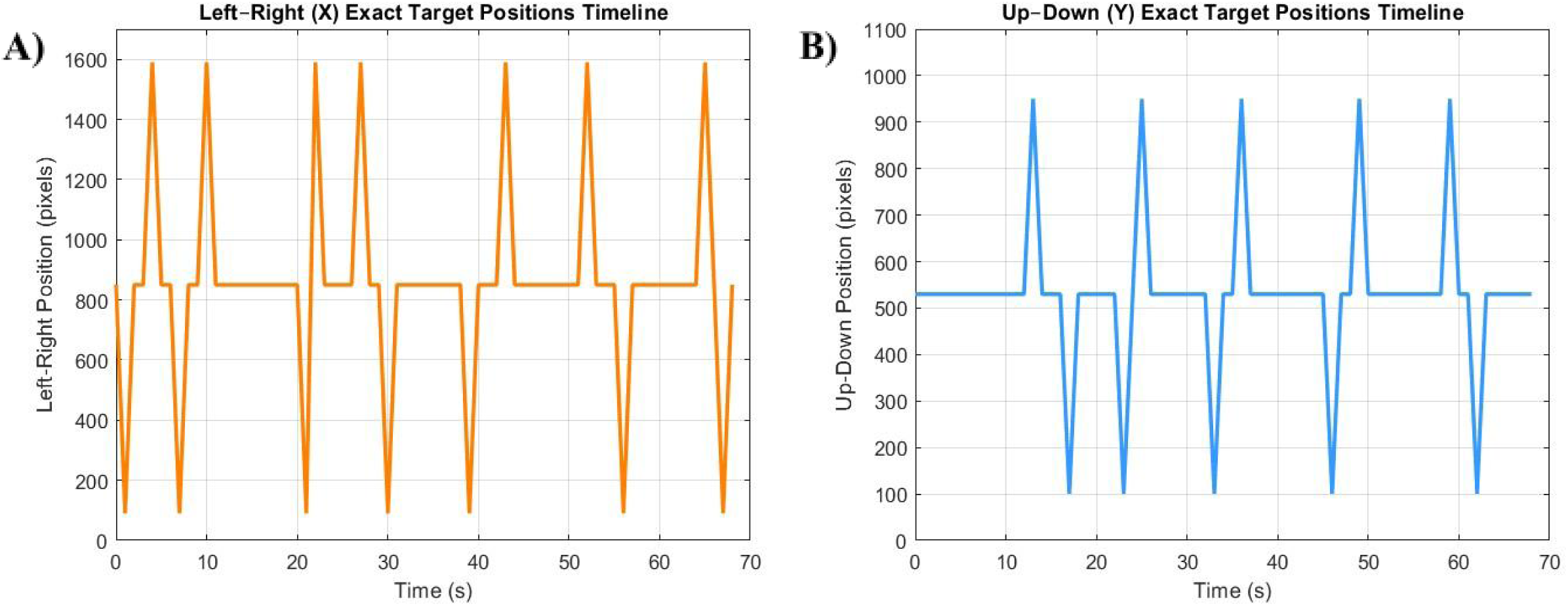
XY Value positions of Training Performance. Exact sequence positions shown in pixels of expected (A) X values and (B) Y values with video/audio timestamp.

**Fig 6.**
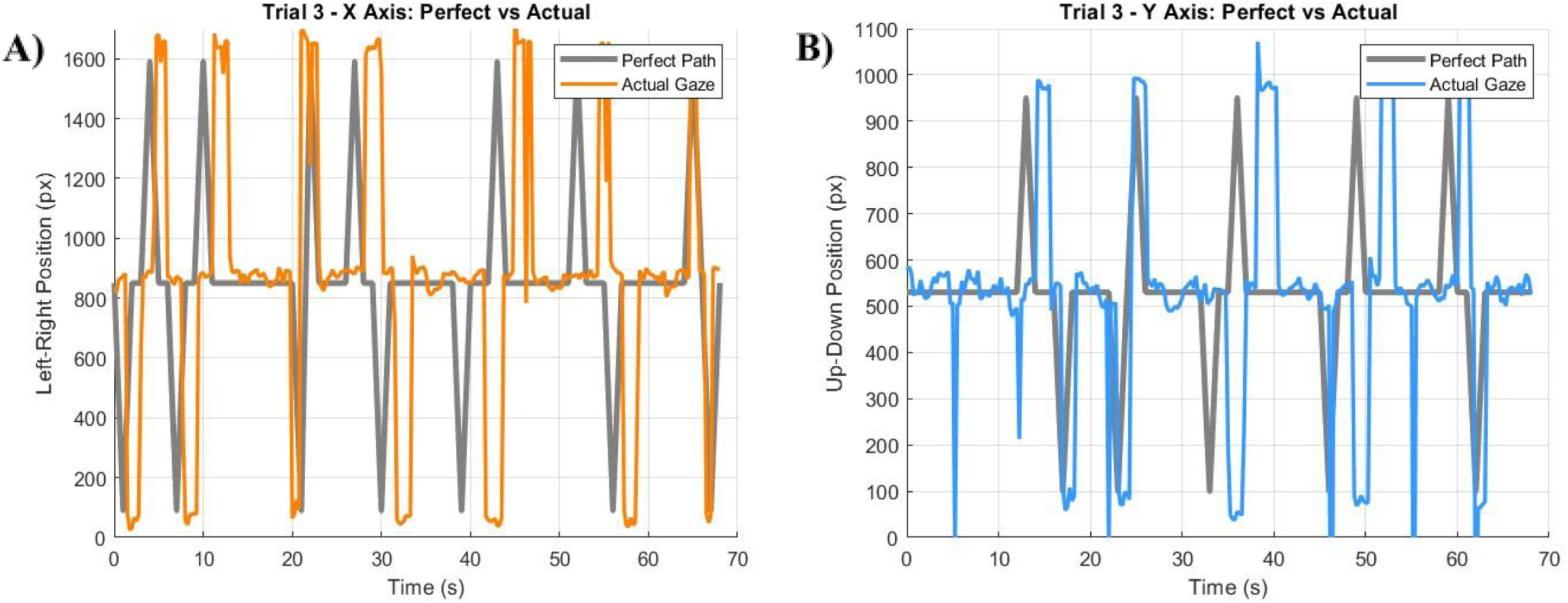
Participant Trial Position and Timing of Actual Performance. Exact sequence positions shown in pixels of individual participant performance in session 5. (A) X values and (B) Y values compared from training position to actual performance with video/audio timestamp

## Results

A repeated-measures ANOVA revealed a significant main effect of sessions on Performance Accuracy, F(4, 36) = 6.99, p < 0.001, an effect size of η²_G_ = 0.26, and Mauchly’s test confirmed that sphericity was met (p = 0.32). An additional repeated-measures ANOVA revealed a significant main effect of sessions on Timing Precision, F(4,36) = 11.67 p < 0.001, an effect size of η²_G_ = 0.25. Mauchly’s test indicated sphericity was violated (p = 0.022); the Greenhouse-Geisser correction confirmed the effect remained significant (p = 0.0086).

Post-hoc pairwise comparisons using Bonferroni correction showed that Session 5 performance accuracy was significantly higher than Session 1 (p = 0.008). No other session pairs differed significantly, though the difference in first and last sessions suggests a gradual increase in performance across each session leading to significant improvement.

Post-hoc pairwise comparisons using Bonferroni correction showed that timing accuracy of movements improved significantly between early and later sessions, with significant differences observed between sessions 1 and 4 (p = 0.044), sessions 1 and 5 (p = 0.0038), sessions 2 and 4 (p = 0.0286), sessions 2 and 5 (p = 0.0055) and sessions 3 and 5 (p = 0.0055). No significant differences were found between later sessions (g4 & g5, p = 1.00), suggesting that timing accuracy plateaued prior to the end of training.

## Discussion

This study investigated the potential of motor sequence learning explicitly using oculomotor function and auditory cues. This novel visual-motor sequence learning task demonstrated significant improvements in both performance accuracy and timing precision, suggesting procedural learning and cognitive-motor coordination can be enhanced through gaze-based repetition. The statistical analyses show a well defined learning curve, proving the efficacy of the task in consolidation of learning following practice. Further analysis of the timing precision can be used to suggest audio cues played a significant role in learning of the sequence. Participants also tended to lag behind audio cue timestamps, in line with findings from Han et al. (2024) and Leow et al. (2025), which suggests auditory stimuli influence motor sequence learning preceding oculomotor movement. Furthermore, it is suggested that audio cues played congruently with visual stimuli encourages stronger development of explicit knowledge within reaction tasks compared to solely visual tasks (Lagarrigue et al., 2021; Silva et al., 2017). The distinct roles of audio and visual stimuli towards influencing eye movement provides a strong foundation for future motor sequence learning studies to consist of neurodegenerative disease or tetraplegic populations, therefore expanding the field on severe motor deficit groups. By using this focused oculomotor approach, the task may potentially aid in designing non-invasive, low-burden rehabilitation tools for enhancing functional outcomes. This gives the opportunity, should interventional and rehabilitational use prove effective, that the task be simplified for frequent/daily use as well.

Various structures of the brain associated with motor-sequence learning and furthermore, the associated oculomotor components of learning, are expected to be pertinent to the results within the current body of work. To elaborate, while motor sequence learning is multifaceted due to various regions of the brain involved, the basal ganglia has been cited throughout prior literature as a core region associated with learning and rewards systems (Di Filippo et al., 2009; Joti et al., 2007). Another region of interest is the ventral tegmental area (VTA), also associated with reward systems (Laviolette & Van Der Kooy, 2001; Morales & Margolis, 2017). These defined areas that encompass the reward system driving learning mechanisms, can be fully traced using fMRI to account for increased functional connectivity by measuring cortical white matter (Hänggi et al., 2010; Tremblay et al., 2021). Furthermore, such increases are noted post-training after motor sequence learning tasks (Jäger et al., 2022), similar results would likely be evident should the present study be replicated with a fMRI component. To further elucidate the mechanisms behind motor sequence learning, neurotransmitter function and measurement must also be accounted for. Once more, neurotransmitter function of motor sequence learning is varied, but namely dopamine (Ferreira et al., 2025), and GABA (Ferreira et al., 2025, Hehl et al., 2025), are promoted by visual tasks, (Tsai et al., 2024), and physical activity (Gao et al., 2025; Petzinger et al., 2013). As elaborated prior, the oculomotor system of the brain has potential to mimic results of physical activity and prompt neurotransmitter signalling. Areas of the oculomotor circuit to include in further analyses include the supplementary motor area (DeSouza et al 2003; Chan, Kucyi & DeSouza, 2015; Jäger et al., 2022; Simon et al., 2024), the frontal eye field (DeSouza et al 2003; Chan, Kucyi & DeSouza, 2015; Rubino et al., 2024), and the supplementary eye field (DeSouza et al 2003), as they are crucial in motor sequence learning (Gonzalez et al., 2016). This novel eye tracking task in future studies, should therefore incorporate a fMRI scanning element to further explore this oculomotor task in having potential results comparable to that of other motor sequence learning tasks.

Throughout our Dance for Parkinson’s work many eye movements are made that are incorporated to gaze to the hand and/or body parts (Bearss et al., 2017). It is made clear that similar results from oculomotor influence can benefit those with neurodegenerative disease and/or those with physical disabilities. Various forms of exercise or dance have been identified previously within the body of work as promoting neuroplastic effects (Dhami et al., 2015). While limited motor movement of the body would likely pose differing results to typical eye-limb motor sequence tasks, the results found in a solely oculomotor task could expand knowledge of an area that lacks specific motor interventions. Parkinson’s disease is an ideal population the novel oculomotor task can be applied, as they experience deficits in areas directly engaged by this learning paradigm (Hamedani et al., 2020). The use of musical cues and reinforcement feedback may promote dopaminergic engagement related to reward circuitry, which is particularly relevant for the PD populations a well (Tsai et al., 2024; D’Allesandro & DeSouza 2025). While not tested within the present study, the exploration of dopaminergic reinforcement involving the score indicator would be of great interest for future studies. Furthermore, dopaminergic changes can modulate anxiety (Baik, 2020; Dong et al., 2020) which future applications of the task can aim to analyze in addition to reinforcement paradigms. Prior studies have discovered physical dance to be cognitively engaging and to promote neuroplastic changes or mitigate future degradation of neural pathways (Barnstaple et al., 2021; Bar & DeSouza, 2016; Cheng et al., 2024; Simon et al., 2024; Quan et al., 2025), thus further supporting the demand for simplified movement related tasks for severe disease cases. Longitudinal studies have urged, (Wong et al., 2018), as understanding mitigative effects of interventions are best imparted when repeated throughout illness progression. Through the implementation of reinforcement paradigms within the task, and engaging areas in which clear deficits are known, there could be potential in promoting greater response or mitigation in neurodegenerative groups such as those with PD.

Within the present study there were several limitations observed. The small sample size of ten participants, while significant, could be increased to get a broader scope of the learning curve and how findings may differ. In addition, a participant expressed an issue with the repetitive nature of the task causing dryness in the eyes, which is to be made note of for future studies involved with eye movement tracking. To understand the full extent of our findings would require fMRI analysis of regions of interest involved in motor sequence learning and furthermore, other pathways exemplifying neuroplastic changes. Additionally, EEG is another measurement tool that could be considered in motor learning applications (Titone et al., 2022). Extra sessions or an extended follow-up session period up to 1 month would be beneficial in a future eye task study to ensure motor sequence learning consolidation and forgetting. A lack of implicit learning is noted, as the explicit learning of the repeated sequence was the central focus of our study. To properly measure implicit learning it is advised to follow learning of the eye dance sequence with a novel sequence. The measurement of saccades is necessary to include for implicit learning (Jiang et al., 2014; Medimorec et al., 2021; Rubino et al., 2025), to assess whether saccadic movements after a novel or random sequence is greater after post sequence training.

These preliminary findings within the current body of work suggest that this novel eye dance task is feasible and sufficient in participant learning of a specific sequence, therefore it can be further investigated in several ways. The simplicity and low physical strain on participants are key components of the eye tracking dance task design and should prove to be cost-effective in training and execution. A psychomotor vigilance task used in the study of passive and active exercise effects on mental fatigue from Jeyarajan et al. (2024), is a task that can be modified under the context of the novel eye tracking task’s results. Its design uses randomized targets appearing on a screen and gauges reaction time measurement by clicking targets, which can be replaced with eye movements and blinks to supplement lack of limb use for future studies. Through the application of our eye tracking test, a future study involving this task could be to account for those with disability or limited limb movement could prove beneficial in promoting neuroplastic effects. Additionally, this would allow for a comparison of other learning paradigms and tasks within the relevant literature to reinforce similarities in sequence learning and elaborate differences regarding the novel task. In doing so first with healthy participants, a clear definition of how such connections develop and adapt over time can be used to compare with a population of those with neurodegenerative diseases.

## Author Contributions

Petrovski, M. - developed task environment, collected participant data, developed code, performed analysis, and first draft of manuscript. Beheiry, S. - collected participant data, edited manuscript. Das, U. U. - created study design and initial task environment. Muhammad, M. - created study design and initial task environment. Simon, R. - conceptualization, funding acquisition Rooprai, S.- conceptualization, initial pilot testing and study design Karimi, A. - conceptualization, initial pilot testing and study design, funding acquisition Bar, R. - conceptualization, funding acquisition DeSouza, J. F. X - conceptualization, initial pilot testing, contributed testing and analysis tools, edited manuscript, and finalized manuscript, funding acquisition.

## Ethics Statement

This study was approved by the Office of Research Ethics Committee (ORE) of York University Certificate #2025-032.

## Conflict of Interest

The authors declare that the research was conducted in the absence of any commercial or financial influence that could be a potential conflict of interest.

## Acknowledgements

The authors would like to thank all the participants that committed their time over the course of several weeks to this study, whose dedication made this possible. We also thank all members of the joeLAB. This research was supported by the Anxiety Research Fund powered by Beneva awarded to York University Faculty of Health.

## Notes

### Competing Interest Statement

The authors have declared no competing interest.

### Summary of Updates

Revised wording of introduction; Added explanation of apparatus and stimuli; Added procedure details; Added statistical analysis subsection; Added additional figures; Revised wording of discussion.

